# Solid-State Nanopore Sizing for cfDNA Sample Quality Control in Point-of-Need Sequencing

**DOI:** 10.1101/2025.03.17.643726

**Authors:** Muhammad Asad Ullah Khalid, Md. Ahasan Ahamed, Anthony J. Politza, Weihua Guan

## Abstract

DNA sequencing is a powerful tool for diagnosing conditions like infectious diseases and cancers. Even though current workflows demand rigorous quality control (QC) of DNA samples, this QC is typically limited to lab settings, despite recent advances in portable nanopore sequencers. For personalized healthcare to truly benefit from the portable sequencer, QC must be performed right where the samples are processed. Here, we present a solid-state nanopore device that provides label-free, controlled quantification and qualification of cell-free DNA (cfDNA). We demonstrated the use of a 1 kbp double-stranded DNA internal marker at a known concentration to measure the concentration of a representative cfDNA target in the presence of genomic DNA. We also found that nanopores with diameters ranging from 6 to 19 nm yield consistent measurements, with a maximum coefficient of variation (CV) of less than 15%. Moreover, analyzing data from multiple nanopores over longer acquisition times can reduce the uncertainty to below 10% CV. Finally, we applied our nanopore QC assay to a plasma cfDNA sample and compared the results with those from a capillary electrophoresis (CE) assay. Both methods produced highly correlated measurements, demonstrating the potential of our nanopore QC assay for effective cfDNA assessment at the point of need.

**Table of Contents Figure:** 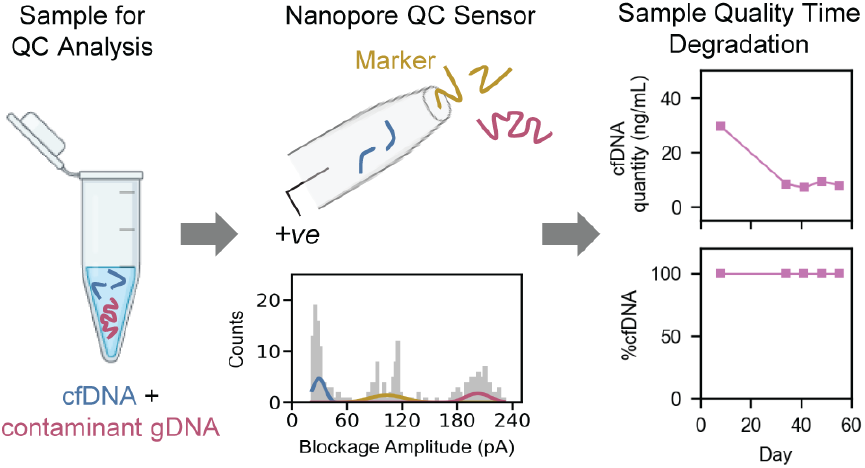

## 1 Introduction

Advancements in DNA sequencing technologies have revolutionized clinical diagnostic and therapeutic practices.^1–4^ However traditional sequencing platforms such as Sanger Sequencing^5^, Illumina Sequencing - Sequence By Synthesis (SBS)^6,7^, Sequence By Ligation (SBL)^8^ and PacBio Single Molecule Real Time (SMRT) sequencing^9^ etc. remain confined to centralized laboratories due to their size, cost, and infrastructure requirements. Point-of-Need sequencing can address these challenges by bringing sequencing capabilities directly to the sample collection site offering rapid, portable and decentralized analysis. It can also enable personalized and faster clinical decision making,^10^ rapid real-time genomic surveillance of infectious outbreaks,^11,12^ and field based environmental and agricultural monitoring^13^. The portable sequencers such as MinION from Oxford Nanopore Technologies (ONT) have demonstrated high potential for enabling (PoN) sequencing to accelerate the developments in personalized medicine for DNA based diagnostics.^14– 16^ However, the genomic sequencing is a multi-step process which include sample collection, sample quality control, and library preparation before subjecting it to actual downstream sequencing analysis.^17^ Although the portability of MinION makes it suitable for PoN applications, the sample quality control remains the bottleneck for true implementation of point-of-need sequencing workflows.

For instance, the quality of cfDNA sample is a critical determinant in sequencing outcomes and should be rigorously assessed before proceeding with downstream library preparation and sequencing workflows. Typically, a larger quantity of total cfDNA and higher relative abundance of mono-nucleosomes in cfDNA samples is ideal for NGS ^18^ mutation analysis towards cancer detection, treatment monitoring and detection of relapses. On the contrary, higher relative abundance of larger sizes (di- and tri nucleosomes) can lead to issues during library preparation steps, potentially causing incomplete adapter ligation or biased amplification.^19^ This can reduce sequencing efficiency and read quality, which also suggests determination of both total cfDNA concentration and % cfDNA before library preparation for sequencing workflow. Additionally, the cfDNA sample can most often be contaminated with high molecular weight (HMW) genomic (g) DNA, from the lysis of leukocytes, released in plasma during sample preparation ^20,21^ leading to decreased sensitivity or inconsistent results in NGS assay.^22^ Similarly, the presence of organic contaminants from extraction processes can easily incur under- or over-estimation of the cfDNA spectrophotometric concentrations.^23,24^ Therefore, tight quality control (QC) steps are required to ensure that extracted cfDNA samples are the right fit for the downstream sequencing workflows. Traditionally, the nanodrop spectrophotometer is used to quantify DNA with limit of detection (LOD) as low as < 1 ng/μL, however it cannot provide information on the fragment length or the DNA integrity of the sample.^25,26^ Similarly, Qubit fluorometric assays can quantify the total dsDNA, ssDNA, or RNA samples but unable to provide fragmentation information of DNA.^27^ To quantify and assess the fragment size profiles of the cfDNA samples, a capillary electrophoresis (CE) is often employed. It performs electrophoretic separation of cfDNA fragments, often categorized as mono-nucleosome (~165 bp), di-nucleosome (~350 bp), and tri-nucleosome (~565 bp) fragments ^28^ because of their apoptotic or necrotic origin in addition to the HMW gDNA (~ 10 kbp or more). However, a typical CE process requires the use of dyes with large input sample quantities, fluorescence/UV detectors, and trained personnel to operate.^29^ Conversely, PCR based approaches focus on the amplification of the specific gene fragment only,^24,30^ which limit their use cases to a wide range of cfDNA targets. Collectively, these approaches are also tedious, time-consuming, and not amendable for field applications. To enable true point-of-need DNA sequencing applications, there is a critical need for a point-of-need amendable sample quality assessment tool.

In this work, we developed a nanopore sensor aiming to perform amplification and label-free quality assessment of plasma cfDNA samples, amendable for point-of-need applications. Our proposed technique utilizes marker DNA as an internal control to perform cfDNA quantification based on size profiling using counts distributions of blockage amplitudes. The multi-gaussian fitting of these blockage amplitude distributions allows the computation of individual event frequencies for the translocation of variably sized DNA components. Subsequently, the event frequencies for the target cfDNA and marker DNA molecules (at known concentration) are then used to estimate the cfDNA concentration. Since the extracted cfDNA samples may contain a fraction of known characteristic HMW gDNA contaminant, our proposed method also enables the relative quantification of total cfDNA (or %cfDNA) as a qualitative metric. We first demonstrated the precise quantification and qualification of 150 bp model DNA fragments as a representative cfDNA target using a 1 kbp DNA marker in mock samples also containing 10 kbp as a representative HMW gDNA at known concentrations. We then analyzed the uncertainty in our nanopore measurements by processing the cumulative data from one, two, three and four nanopores with data acquisition times from 5 to 30 min for a single mock sample with fixed concentrations of individual model fragments. Our findings revealed that the measurement uncertainty decreased with longer data acquisition times but increased with the number of nanopores used. We further evaluated the applicability of nanopore QC assay by analyzing the aging of a commercially purchased plasma sample of a healthy individual. Both the nanopore QC assay and a traditional capillary electrophoresis assay showed similar degradation trends in measured cfDNA concentration, with no effect on %cfDNA. The comparison between our nanopore QC assay and the capillary electrophoresis assay indicated negligible differences in individual measurements of cfDNA concentration and %cfDNA. These results suggest that the nanopore QC assay is a robust and comprehensive tool for cfDNA sample quality assessment.

## 2 Results and Discussion

### 2.1 cfDNA sample quality control using nanopore size counting

To assess the quality of cfDNA in complex human plasma or serum samples, a typical sample preparation workflow has been shown schematically in **Figure 1a**, where blood is collected from healthy or diseased individuals in EDTA collection tubes followed by two step centrifugation at 2000 × g at 4 °C for 10 min for plasma separation and aliquots preparation.^20,31^ These plasma samples are then used for cfDNA extraction using, for example, commercially available silica-based membrane columns ^32^ according to the manufacturer’s protocol. Before any downstream analysis, these extracted samples are subjected to Agilent Tapestation 4150 or Bioanalyzer 2100 (for CE), Nanodrop spectrophotometer (Thermo Fisher Scientific, Waltham, MA) or Qubit fluorometer (Invitrogen, Life Technologies) for size profiling and quantitative analysis respectively.^24^ We present here a glass nanopore sensor for the comprehensive quality assessment of these extracted cfDNA samples as shown in **Figure 1b** due to its ability to perform label-free size counting of short fragments. The extracted cfDNA samples are diluted with a 1 kbp dsDNA marker having 100 pM final concentration in a 4 M LiCl Tris-EDTA (pH 8.0) buffer as an internal control. The use of high salt concentration as a nanopore measurement buffer can be signified by its detection capability of short DNA fragments^33^ without any requirements of labels, nanopore surface modifications or use of hydrogels as entropic barriers to slow down DNA translocations. A ~10 nm nanopore diameter fabricated using a laser puller, filled with the salt solution, is inserted in the complex DNA sample containing cfDNA. A positive bias is applied on the patch electrode for ionic current data recording. By analyzing a ~ 25 – 30 min *I-t* trace data of the nanopore using a custom MATLAB script, the event scatter is plotted to obtain counts distributions of blockage amplitudes. A multi-gaussian fit to the blockage amplitude distributions allows for the computation of individual event frequencies for the translocations of cfDNA targets, marker (M) DNA (1 kbp) and HMW gDNA. To measure the total concentration of cfDNA an internal calibrator assisted concentration measurement approach is adapted from Charron et al.^34^ The precisely known concentration of our internal 1 kbp marker (C_M_) along with cfDNA target and marker frequencies (*f*_*T*_ and *f*_*M*_) are used to mathematically estimate the total cfDNA concentration (C_T_) using 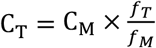 relation valid for diffusion limited transport. Similarly, the cfDNA target frequency (*f*_*T*_) and the HMW gDNA frequency (*f*_*gDNA*_) values are used for %cfDNA measurement using %cfDNA 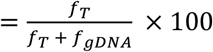. The frequency values are used here instead of total counts to compensate for data loss due to possible timed clogging of the nanopores during experiments. The estimated C_T_ and %cfDNA values are then used as QC metrics of cfDNA for decision-making towards NGS library preparation. A multiparametric comparison of proposed technology with standard laboratory procedures has been presented in **Figure 1c**.

**Figure 1.**
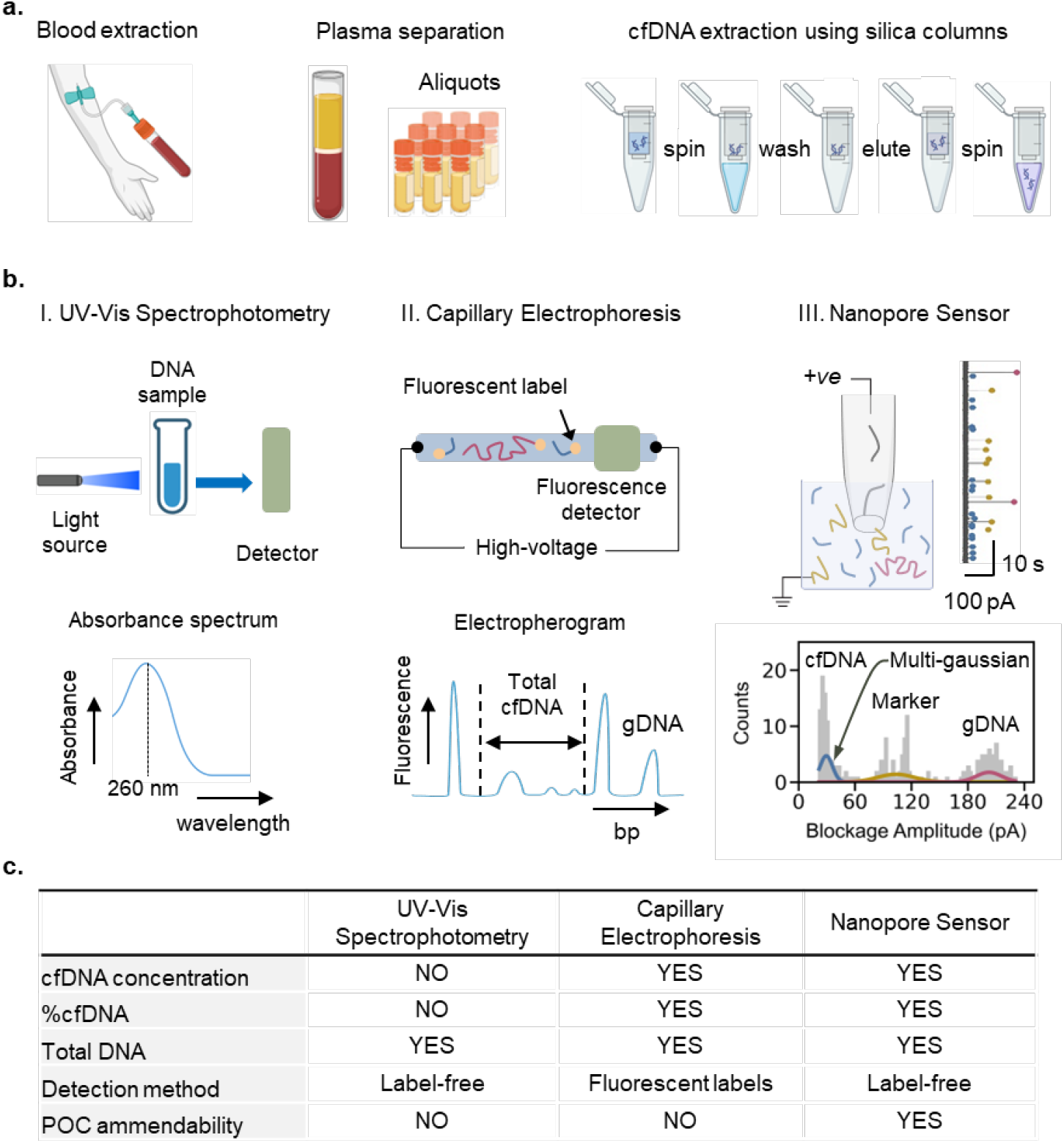
Scheme for cfDNA sample quality control using conventional strategies and proposed nanopore sensing: **(a)** Blood collection for plasma separation by double centrifugation with aliquots preparation followed by cfDNA extraction using silica-based membrane columns according to manufacturer’s protocol before subjecting to quality control procedures. **(b)** Conventional cfDNA quality control procedures include I. UV-vis spectrophotometric absorbance at 260 nm which only provides total DNA concentration, and II. Capillary Electrophoresis which can provide total DNA quantity, total cfDNA quantity and %cfDNA but requires fluorescent labels, detector and high voltage operation. However, our proposed label free glass nanopore sensor scheme performs quantification of extracted total cfDNA as well as %cfDNA from the complex DNA samples containing 1 kbp DNA marker. This is accomplished by processing ionic current - time data for single molecule analysis of the complex DNA sample using blockage amplitude distribution and multi-gaussian fitting to extract individual frequencies *f*_*T*_, *f*_*M*_, and *f*_*gDNA*_ of the component cfDNA, marker and HMW gDNA fragments. The frequencies are then used for the estimation of total cfDNA concentration and the determination of %cfDNA. **(c)** A multi-parametric comparison of different quality control techniques.

### 2.2 Quantification and qualification of 150 bp dsDNA from the mock samples using nanopore sensor

To enable the quantification of cfDNA in the plasma samples using nanopore sensor, we first evaluated the ability of our nanopore sensor to quantify 150 bp dsDNA as a representative cfDNA target (T) in a mock sample with 10 kbp dsDNA as high molecular weight (HMW) gDNA contaminant, and 1 kbp dsDNA as internal marker (M) in 4 M LiCl Tris-EDTA (pH 8.0) measurement buffer. Seven different mock samples were prepared by varying the concentration of 150 bp target in the range of 10 pM to 200 pM and keeping the concentrations of marker and HMW DNA constant at 100 pM each. These test concentration ranges were carefully chosen to reflect the coverage of probable yields of cfDNA in the actual plasma which were to be eluted and then diluted in the 4 M LiCl measurement buffer. All the mock samples were tested using glass nanopore devices fabricated using laser pipette puller with diameters between ~ 8 to 18 nm. The ionic current-time (I-t) data were recorded at 100 kHz sampling frequency and 5 kHz filter for all the mock samples using Axon Axopatch 200B amplifier for 30 min each as single measurement. The current blockage event data was then extracted using a custom MATLAB script for each mock sample. This data was further processed in a custom python program to plot counts distributions of blockage amplitudes with multi-gaussian fitting using a Gaussian Mixture Model (GMM). The total individual counts under each gaussian were then used to determine the individual event frequencies of 150, 1k and 10k bp dsDNA targets and hence the concentration and % of 150 bp dsDNA as previously discussed. In **Figure 2a**, the representative *I-t* traces of seven different mock samples have been shown. These I-t traces implicitly show the increasing frequency of the translocation events for 150 bp dsDNA as indicated by blue circles in a concentration dependent manner. This was further confirmed by the associated muti-gaussian fittings of the counts’ distributions of blockage amplitudes for all seven mock samples as shown in **Figure 2a**, where the relative counts for 150 bp dsDNA translocations were increasing. The gaussian distributions for 1k and 10k bp dsDNA show successively decreasing relative counts distributions despite their fixed concentrations in the mock samples. This is understandable because the capture in diffusion limited transport is DNA length independent while simultaneously being concentration dependent.^34^ An above 99% correlation between measured and actual concentrations and % of 150 bp dsDNA as shown in **Figure 2b and c** suggested the highly precise quantification ability of our nanopore sensor in mock samples containing three different targets with length ratios as large as 67×. Although data for the various mock samples were obtained using nanopores with diameters ranging from 8 to 18 nm, a systematic analysis was subsequently conducted to examine the nanopore size-dependent variations in the measurements of concentration and % of 150 bp dsDNA as discussed below.

**Figure 2.**
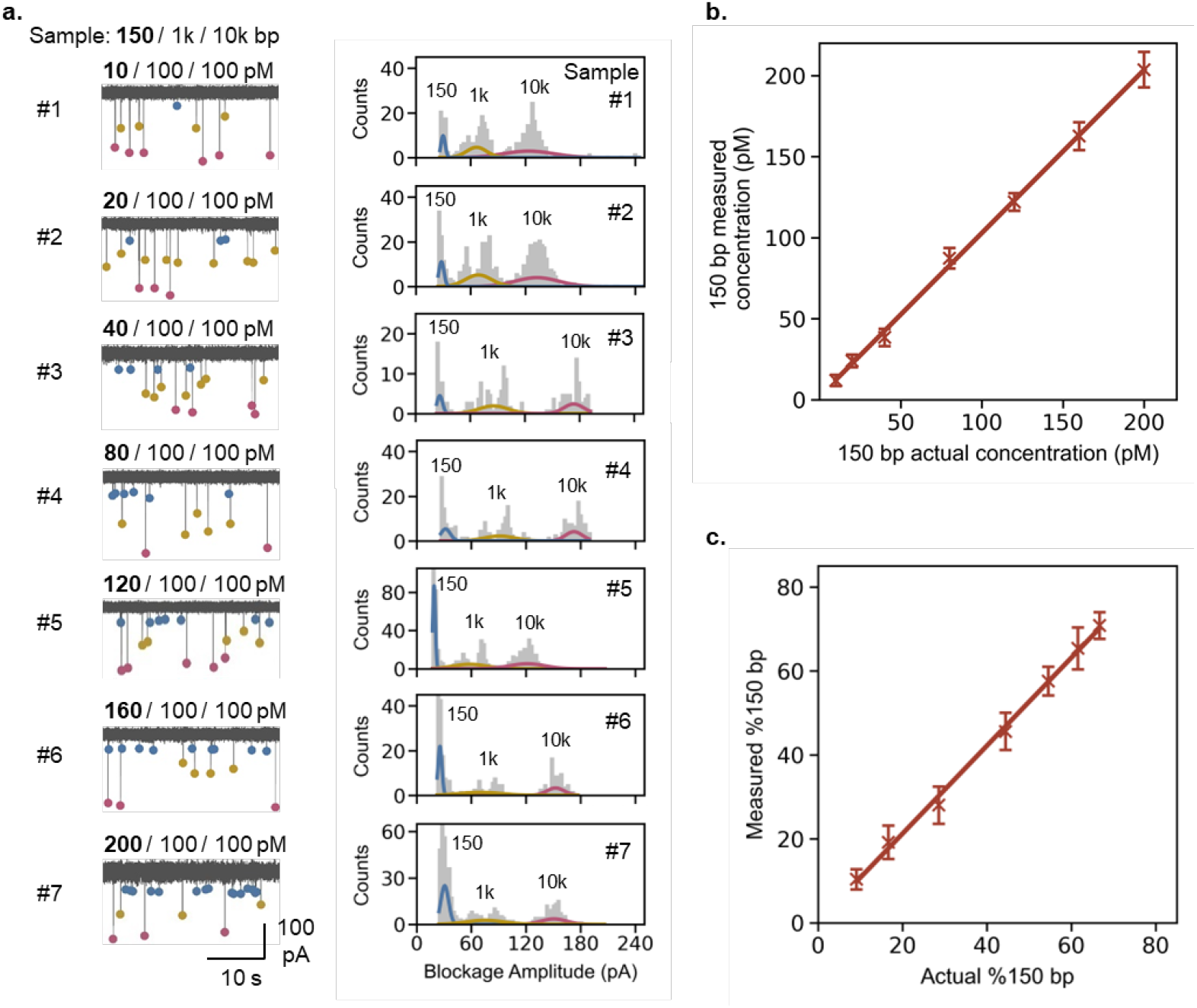
Nanopore sensor-based quantification and qualification of 150 bp dsDNA from different mock samples. **(a)** Representative I-t traces and counts histograms of the mock samples with variable concentrations (10 – 200 pM) of 150 bp dsDNA, where concentrations of 1 kbp marker (M) dsDNA and 10 kbp dsDNA (HMW) were fixed at 100 pM each. Correlation of measured vs. actual **(b)** concentration (pM) and **(c)** % of 150 bp dsDNA. Data has been presented as μ ± σ (n = 3).

To deal with the heterogeneity of the nanopore size for the measurements, four different glass nanopores were fabricated using laser pipette puller with respective pore diameters estimated to be ~ 6.2, 9.7, 12.2 and 18.9 nm. The IV curves and open pore conductance values used for pore size estimations have been provided in **Figure S1.1**. Each glass nanopore was tested with the mock sample of 150, 1k and 10k bp dsDNA with fixed concentrations of 120 pM, 100 pM and 100 pM respectively. The capture rate is expected to increase with the increase in nanopore diameter (from 6.2 to 18.9 nm) and hence the capture radius which can be seen from the *I-t* traces and multi-gaussian fittings of the counts distributions of blockage amplitudes in **Figure 3a**. The increase in nanopore diameters can also be confirmed from the shrinkage of blockage distributions of 1k and 10k bp, for Np # 1 to 4, on the blockage amplitude scales. Interestingly these populations also imply relatively higher counts for 150 bp dsDNA at 120 pM as compared to 1k and 10k bp at 100 pM each which is expected. The concentration and % of 150 bp dsDNA were estimated using a previously established method and plotted against nanopore diameter (Np *ϕ*) (data presented as n = 3, μ ± σ) as shown in **Figure 3b and c**. The dashed lines show the actual concentration of 120 pM and 54.5% of 150 bp dsDNA respectively. The coefficient of variation (CV) for measured 150 bp concentration of Np #1 to 4 were found to be 11.9, 5.1, 7.3 and 14.4 % respectively suggesting negligible variability between concentration measurements. Similarly, the CV values of 8.6, 5.8, 10.3 and 10.7% for measured %150 bp also implied insignificant variability of these measurements. This suggested that the nanopore diameter did not affect the measurements significantly and hence enabling the use of our nanopore sensor for QC of actual %150 bp from plasma samples. As the analysis time is dependent on the time for data acquisition, we further sought to evaluate the cumulative data from multiple nanopores and for different data acquisition times before the validation studies using plasma cfDNA sample.

**Figure 3.**
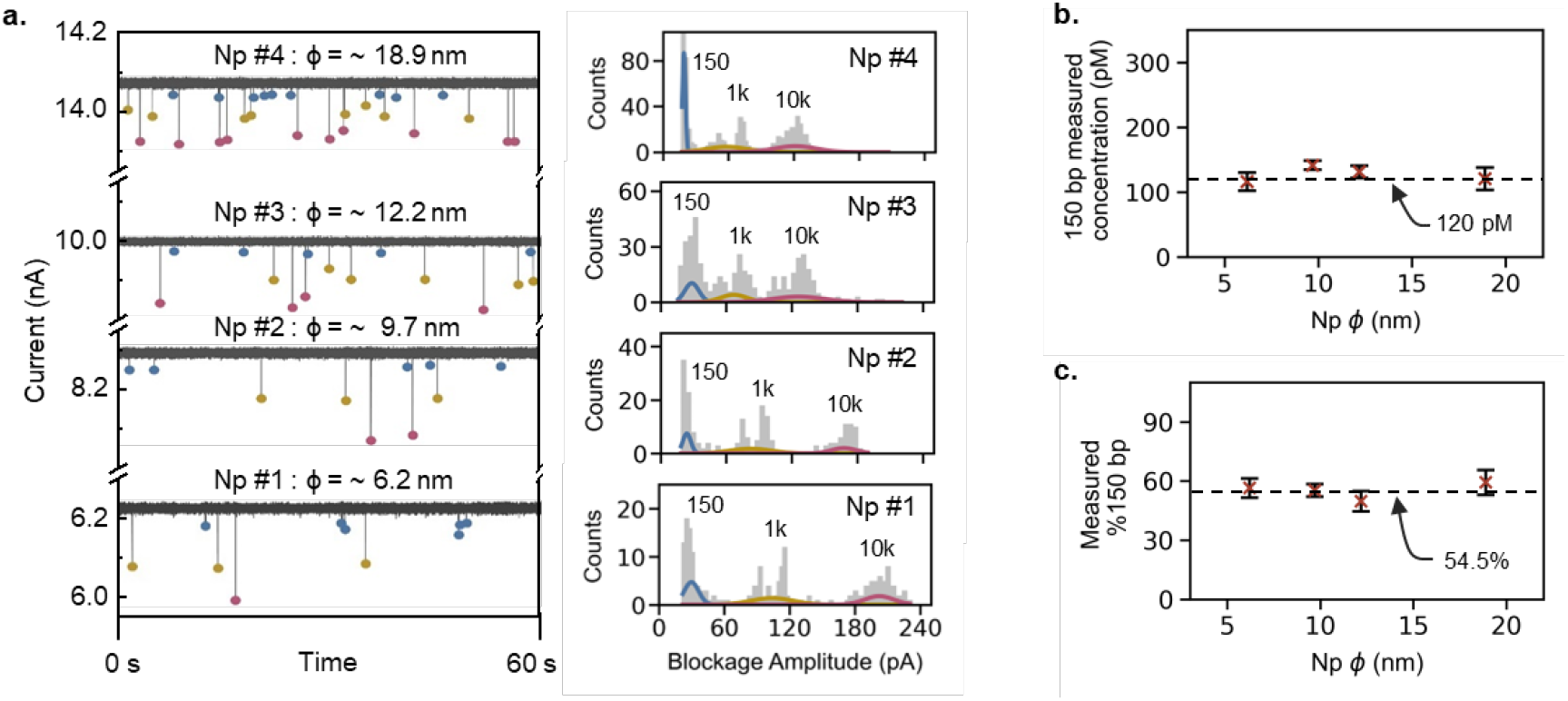
Quantification and qualification of 150 bp dsDNA from a fixed mock sample using different nanopore sensors. **(a)** Representative I-t traces and counts histograms for the mock sample 150 / 1k / 10k bp (at 120 / 100 / 100 pM) obtained using different nanopores with diameters in the range of 6-19 nm. **(b)** Measured concentration of 150 bp and **(c)** measured %150 bp vs. Np *ϕ* (nm). Data has been presented as μ ± σ (n = 3).

### 2.3 Evaluating measurement uncertainty of nanopore QC assay

To assess the uncertainty of our nanopore sensor measurements, we analyzed the cumulative data from multiple nanopore configurations with different data acquisition times. The nanopore configurations of one, two, three, and four nanopores were compared against data acquisition times of 5, 10, 15, 20, 25, and 30 each creating a total of twenty-four test combinations. The mock sample of 150 : 1k : 10k bp at fixed concentrations of 120 : 100 : 100 pM respectively was tested to compute the measurement errors in all the twenty-four test combinations. Four different nanopores were fabricated using laser pipette puller with diameters estimated within 9.7 ± 0.8 nm using open pore conductance data from IV characteristics shown in **Figure S1.2**. Each nanopore was used to acquire current-time (I-t) data in 5 min chunks for a total of 90 min using Axopatch 200B amplifier system which was processed using a custom MATLAB script to obtain blockage amplitudes as described previously. The processed data was then divided into twenty-four test combinations as described above according to the number of pores and data acquisition times. The measurements of concentration and % of 150 bp for each combination were performed according to the internal marker (1 kbp) controlled method described previously. The representative counts distributions of blockage amplitudes along with their respective multi-gaussian fittings have been presented in **Figure S1.3**. The measurement uncertainty was determined in terms of %CV. A heatmap of the uncertainties in measured concentrations of 150 bp dsDNA has been presented in **Figure 4a**. The measurement uncertainty or %CV tends to decrease as the time of data acquisition and number of nanopores increase, which is expected as the event count also increases.^35^ Subsequently, the % CV values for measured %150 bp shown in the heatmap in **Figure 4b** also follow the similar trend. These results indicate that measurement uncertainties can still be less than 15% for the measurement of 150 bp concentration and %150 bp if the data acquisition time is reduced to 15 min for larger number of nanopores with insignificant size variations. A CV of 15% is typically acceptable for the lab-based capillary electrophoresis systems. Given its suitability for POC settings, a multiple nanopores strategy with parallel data acquisition capability can significantly reduce the turn-around time of the nanopore QC assay.

**Figure 4.**
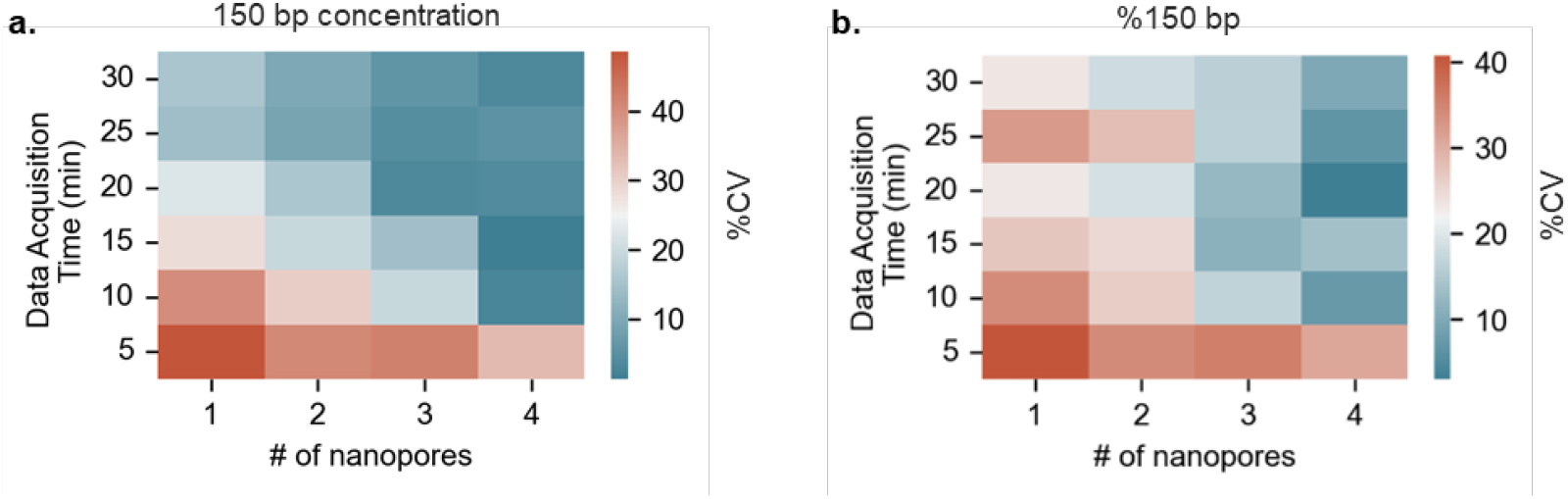
Utilizing data from multiple nanopores to potentially reduce the measurement error and turnaround times of the nanopore QC test. Heatmaps for %CV values of measured **(a)** 150 bp concentrations and **(b)** %150 bp from different combinations of data acquisition times and # of nanopores (number of measurements n = 3, μ ± σ).

### 2.4 Validation of nanopore QC assay using plasma cfDNA sample

To validate our nanopore QC assay for the quantification and qualification of cfDNA, a commercially purchased plasma sample of a healthy individual was subjected to our nanopore QC and gold standard TapeStation capillary electrophoresis (CE) assays. The time degradation study of the cfDNA was performed for nearly seven weeks of extractions after plasma separation post blood collection. The 1 mL aliquots of unused plasma were stored at −80 °C before use. The cfDNA from the 1 mL plasma aliquot was extracted in 55 μL of elution buffer each week using QIAamp circulating nucleic acid kit according to the manufacturer protocol. The extraction process has been detailed in subsection 4.3 of the “Materials and Methods”. The eluate was directly subjected to TapeStation characterization. However, it was further diluted in the 4 M LiCl salt buffer by a factor of 40× for nanopore QC assay. We chose a factor of 40× dilution to allow the plasma concentrations fall in the range of 2 – 40 ng/mL which will then correspond to the previously established detection range of 10 – 200 pM for 150 bp dsDNA. The details about the TapeStation CE assay have also been presented in subsection 4.4 of the “Materials and Methods”. The amount of cfDNA and %cfDNA for quantification and qualification were obtained from capillary electropherograms which have been shown in **Figure 5a** on the left side for respective day of cfDNA extraction post-plasma separation. To calculate the concentration of cfDNA and %cfDNA from electropherograms, the area under the curve between 50 – 600 bp markings is determined and compared with the total coverage area. The known concentrations of lower and upper markers then allow for the computation of cfDNA concentration and %cfDNA in the eluate which were then back calculated to the original plasma concentrations considering a 100% recovery of cfDNA from 1 mL of plasma to 55 μL of eluate. Each extracted cfDNA was also concurrently subjected to our established nanopore QC assay after dilution with the 4 M LiCl measurement buffer. The nanopore QC data of each respective day of cfDNA extraction has been presented as the normalized counts’ distributions of blockage amplitudes with their corresponding multi-gaussian fittings in **Figure 5a** on the right side. A quick analysis of the data from both the TapStation CE and nanopore QC assays suggests that no HMW gDNA was detected which is probably due to the collection of blood in streck tubes. Streck tubes are specifically designed to stabilize cfDNA and inhibit cellular degradation or hemolysis which may cause gDNA contamination. The absence of gDNA characteristic peaks in both the TapeStation and CE assays implied that a %cfDNA was 100% and remained unaffected throughout the seven weeks of degradation study as shown in **Figure S1.4**. The measured concentrations of the cfDNA from each respective extraction have been plotted as shown in **Figure 5b** for both TapeStation CE and nanopore QC assays. The concentration of cfDNA significantly dropped from the first week extraction to fourth week extraction after plasma separation from blood and leveled off below 10 ng/mL. This suggests that a plasma sample stored for longer than one week after separation from blood rapidly loses the quality of cfDNA in terms of the measured quantities. Despite significantly lower input cfDNA abundance a concentration above 2 ng/mL of cfDNA in the plasma sample has still been reported for the NGS library preparation protocols.^36^ Generally, multiple extractions can be run or use of smaller elution buffer volumes can increase the cfDNA yield for improved coverage in sequencing workflows for healthy control samples. The concentration of cfDNA and %cfDNA are specific to each individual but the availability of a point of need sample QC tool can significantly add value to decision making for NGS library preparation especially when the sample degradation timeline is very short. A high correlation was observed for the measured cfDNA concentration between nanopore QC and TapeStation CE assays as shown in **Figure 5c**. Our nanopore sensor is a label-free, electronic readout approach for sample QC with performance comparable to the conventional laboratory based TapeStation CE assay which suggests that it has a high potential for POC implementation to complement modern-day needs of portable nanopore sequencers.

**Figure 5.**
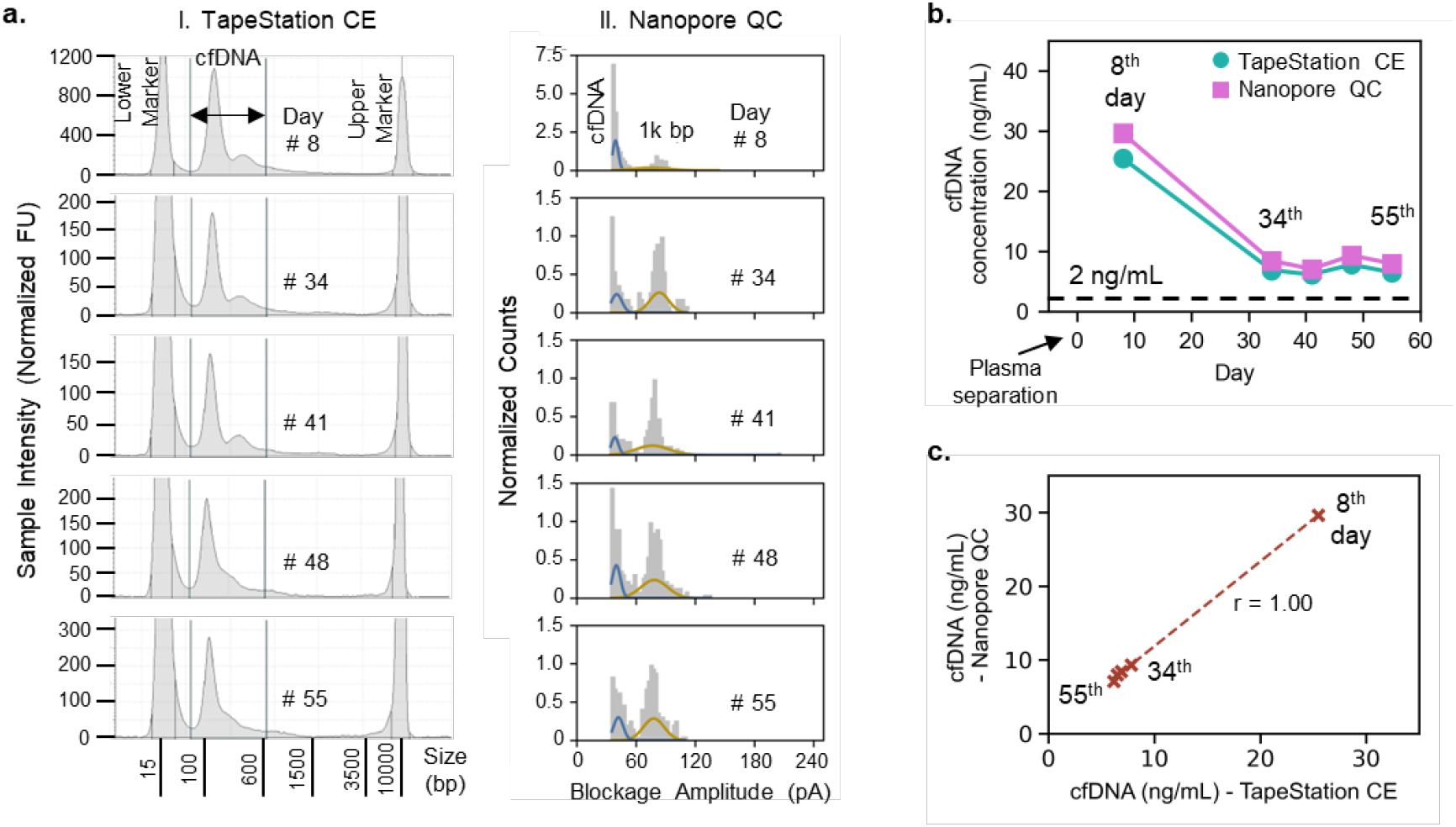
Validation of nanopore QC test using the plasma sample of a healthy individual. **(a)** Electropherograms obtained from TapeStation capillary electrophoresis (CE) assay with corresponding counts distributions of blockage amplitudes from nanopore QC assay. **(b)** Consistent drop observed in measured cfDNA concentrations using both TapeStation CE and nanopore QC assays at different days up till 55^th^ day post-plasma separation. cfDNA concentration. **(c)** Measured concentrations of cfDNA using both Nanopore QC and TapeStation CE assays showed high correlation validating the proposed assay’s QC performance.

## 3 Conclusion

With the rapid miniaturization of sequencing technologies to enable point-of-care (POC) diagnostics for personalized healthcare, there is an urgent need for a comprehensive POC sample quality control (QC) tool. Here, we present a nanopore sensor for the assessment of cfDNA samples, amenable for point-of-need applications. By employing a 1 kb DNA marker at known concentration, our technique precisely quantifies the varying concentrations of a 150 bp dsDNA as representative cfDNA target in a test range of 10 – 200 pM in mock samples, as a quantification metric in QC assay. Additionally, it enabled the precise measurement of %150 bp as a qualification metric of sample QC in the presence of a 10 kbp dsDNA a representative HMW gDNA contaminant. The estimated and known values of concentration and % of 150 bp dsDNA showed a high correlation of above 90%. Smaller variations in nanopore diameters (6 – 19 nm) minimally affected the measurements with a CV of <15%. We have further demonstrated that the measurement uncertainty can be tuned by analyzing the cumulative data from multiple nanopores for different data acquisition times. The measurement uncertainty decreases as the number of nanopores, and the data acquisition times increase. Validation studies, tracking cfDNA degradation over nearly seven weeks using our nanopore sensor and TapeStation CE assay, showed strong correlation in concentration measurements with significant cfDNA degradation observed after four weeks of plasma separation post blood collection. Given the label-free simple electronic readout capabilities, this method holds high promise as a POC amendable sample QC tool for point-of-need NGS.

## 4 Materials and methods

### 4.1 Materials and chemicals

For nanopore fabrication, quartz capillaries with inner and outer diameters of 0.5 and 1 mm respectively (Q100-50-7.5) were purchased from Sutter Instrument, USA. A microinjector for filling the nanopipettes (MF34G-5) was purchased from World Precision Instruments. The nanopipette holder (QSW-T10N) and 0.2 mm diameter Ag wires were purchased from Warner Instruments, USA. UltraPure™ DNase/RNase-Free Distilled Water (Catalog number: 10977015) and the dsDNA fragments of various lengths (150, 1k, and 10k bp) were purchased from Thermofisher. Tris-EDTA-buffer solution (pH 8.0), lithium chloride (LiCl) salt, sulfuric acid (H_2_SO_4_) and hydrogen peroxide (H_2_O_2_) were purchased from Sigma-Aldrich. QIAamp circulating nucleic acid kit (catalogue number: 55114) was purchased from QIAGEN for cfDNA extraction from plasma samples. A plasma sample of a healthy control was purchased from BioCollections Worldwide, Inc. High Sensitivity D5000 (HSD5000) ScreenTape and Reagents (part numbers: 5067-5592 and 5067-5593) were purchased from Agilent Technologies.

### 4.2 Nanopore fabrication

To fabricate the glass nanopores, glass capillaries were cleaned using piranha solution which was prepared in lab by mixing H_2_SO_4_ and H_2_O_2_ in 3:1. Briefly, the capillaries immersed in piranha solution were placed on a hot plate for 30 min at 85 °C followed by rinsing with DI water and vacuum drying at 120 °C for 20 min. The capillaries were then subjected to a two-line recipe in laser pipette puller (P-2000, Sutter Instruments, USA) to fabricate nanopores. Line 1: “heat 750, filament 5, velocity 50, delay 140, and pull 50”, Line 2: “heat 715, filament 4, velocity 30, delay 145, and pull 215”. This recipe fabricates nanopores with diameters typically around 10 nm. However, to change the nanopore diameters, we slightly adjusted the “pull” parameter in line 1 and 2. The nanopore diameters were estimated from open pore conductance values obtained through IV characteristics as described in previous work.^35^ The pulling parameters of fabrication recipe are instrument-specific, and the process is sensitive to the physical conditions of the environment, so the fabrication recipe can be customized to obtain desired nanopore diameters.

### 4.3 Extraction and purification of cfDNA from plasma samples

Various pre-analytical factors may affect the total cfDNA recovery,^37^ so an extensively adopted plasma processing and extraction protocol was employed. Briefly, the fresh plasma sample was obtained 2 h post blood collection from healthy individual (with no HIV and cancer history) following double centrifugation procedure with custom protocol by the vendor BioCollections Worldwide, Inc and stored immediately at −80 °C. The plasma sample was received frozen on day 7^th^ post-plasma separation and stored at −80 °C before cfDNA extraction. The QIAamp circulating nucleic acid kit has been shown to recover >90% of the total cfDNA post-extraction. The extraction procedure was performed according to the manufacturer’s protocol which involved four typical steps of silica column extractions; lyse, bind, wash and elute. The 1 mL of plasma was used with an elution volume of 50 μL. And the eluted volume was then further used for qubit, capillary electrophoresis (CE) and nanopore cfDNA QC.

### 4.4 Agilent TapeStation CE assay for cfDNA size profiling and %cfDNA

Following the cfDNA extraction from plasma samples of healthy controls, the eluted cfDNA samples are subjected to a HSD5000 ScreenTape Assay in Agilent TapeStation 4150 system for size profiling and % cfDNA evaluation. The HSDD5000 reagents (ladder and sample buffer) are brought to room temperature for 30 min. The HSD5000 ScreenTape device is inserted into the ScreenTape nest of the 4150 TapeStation instrument. Appropriate selections of required sample positions are made in the TapeStation Controller software. Reagents and samples are vortexed and spun down before use. To prepare the ladder, 2 μL of HSDD5000 sample buffer and 2 μL of the ladder are added at position A1 in a tube strip. Whereas, for each sample 2 μL of HSD5000 sample buffer and 2 μL of cfDNA sample are added at the subsequent positions in the tube strip. The tube strip is then capped and then liquids are mixed using a vortex mixer at 2000 rpm for 1 min followed by spinning down for 1 min. The tube strip is also loaded in the 4150 TapeStation instrument by confirming the A1 position of ladder on tube strip holder. The cap of the tube strip is carefully removed ensuring all the sample volume is sitting down at the bottom. The instrument is run and the TapeStation analysis software opens automatically afterwards to display the results.

### 4.5 Data analysis method and statistics

All ionic current-time (I-t) data were acquired at 100 kHz sampling frequency using a patch clamp amplifier Axopatch 200B by Molecular Devices, an NI 6363 DAQ card, and a low pass filter (5 kHz) in a custom LabVIEW program. Data was further analyzed using custom MATLAB and Python scripts to extract peak information, and process for multi-gaussian fittings respectively. Python script for obtaining counts histograms and multi-gaussian fittings has been provided in **S2. Supplementary Code**. All the measurements were repeated at least three times, unless mentioned otherwise.

## 5 Acknowledgments

This work was partially supported by the National Science Foundation (2319913, 2045169) and the National Institute of Health (R33AI147419), and USDA (NIFA 2022-11225). Any opinions, findings, conclusions, or recommendations expressed in this work are those of the authors and do not necessarily reflect the views of the National Science Foundation, the National Institutes of Health, or the USDA.

## Notes

### Competing Interest Statement

The authors have declared no competing interest.

